# Mechanisms analysis for Formononetin counteracted-Osimertinib resistance in Non-small-cell lung cancer cells: from the insight into the gene transcriptional level

**DOI:** 10.1101/2023.09.27.559871

**Authors:** Runyang Yin, Jing Gao, Yang Liu

## Abstract

**Backgrounds:** Osimertinib resistance in Non-Small-Cell Lung Cancers (NSCLC) is a difficult problem to be solved in clinic. Formononetin is one of the main isoflavone components, which has strong anti-cancer effects in NSCLC. However, the potential effects and the mechanisms of Formononetin to counteract the Osimertinib resistant in NSCLC is remain to be uncovered.

**Methods:** Osimertinib-resistant NSCLC cell line (H1975_OR) was generated by increasingly adding of Osimertinib in H1975 cell culture medium. The Formononetin was used to induce H1975_OR cell apoptosis an to inhibit cell proliferation and clonal formation. The RNA-sequencing was used to study the potential mechanisms for Formononetin to counteract the Osimertinib resistant in NSCLC.

**Results:** Formononetin could significantly induced cell apoptosis, whereas, dramatically inhibited cell proliferation and clinal formation in H1975_OR cells. Formononetin induced tremendous alterations of gene expression in H1975_OR cells and these genes can be clustered into unique GO, KEGG and GSEA pathways.

**Conclusion:** Our results preliminarily uncovered the underlying mechanisms for Formononetin counteracted Osimertinib-resistance in NSCLC cells and provided a potential treatment method for Osimertinib resistant NSCLC patients.

## 1 Introduction

Lung cancer is a malignant tumor originating from lung epithelial tissue, which has the highest incidence and mortality in China and even in the world ^[1]^. Lung cancer has two major subsets, namely non-small cell lung cancer (NSCLC) and small cell lung cancer (SCLC). NSCLC is a heterogeneous tumor that accounts for approximately 85% of all newly diagnosed lung cancers ^[2]^. The incidence and mortality of NSCLC are rising rapidly, posing a serious threat to human life and health and putting great pressure on the medical system. Therefore, the in-depth study on the pathogenesis and treatment strategy of NSCLC has far-reaching significance.

Over the past decade, in-depth analysis of lung cancer genomes and signaling pathways has further defined NSCLC as a unique group of disease with genetic and cellular heterogeneity ^[3]^, in which, smoking, radiation exposure and air pollution were clustered as the major risk factors for the disease ^[4]^. Most patients are diagnosed with advanced disease due to inadequate screening procedures and late onset of clinical symptoms. As a result, the patient’s prognosis is usually poor ^[5]^. Several diagnostic methods are available for NSCLC, including X-ray, Computed Tomography (CT) and Positron Emission Tomography (PET) imaging, and histological examination of tumor biopsies ^[6]^. To determine the best treatment strategies, accurate staging of the cancer is required, including surgery, radiochemotherapy and immunotherapy, and targeted therapies such as anti-angiogenic monoclonal antibodies, Tyrosine Kinase Inhibitors (TKIs), or immunotherapy if the tumor carries an oncogene mutation ^[7, 8]^。

Osimertinib is an irreversible third-generation Epidermal Growth Factor Receptor (EGFR) Tyrosine Kinase Inhibitors (EGFR-TKIs) that simultaneously inhibits both EGFR-sensitive mutations and EGFR-T790M resistant mutations ^[9]^. Compared with first-generation and second-generation EGFR-TKIs, Osimertinib has a stronger ability to penetrate the blood-brain barrier and has good efficacy in patients with central nervous system metastases ^[10]^. In November 2015, the US Food and Drug Administration (FDA) approved Osimertinib for the treatment of patients with EGFR-T790M mutation in NSCLC. It has also become the fastest anti-cancer drug ever launched by the US FDA: from clinical to market in only 2 years ^[11]^. In March 2017, Osimertinib was first approved in China for use in adult patients with locally advanced or metastatic NSCLC who developed disease during or after treatment with EGFR-TKIs and tested positive for EGFR-T790M mutation ^[12]^. In December 2020, based on data from the ADAURA, Osimertinib was approved in the US for the adjuvant treatment after tumor resection in adult patients with EGFR exon 19 deletion or exon 21 L858R mutation in NSCLC ^[13]^.

Previous data told us that the median progression-free survival (mPFS) was 18.9 months in the Osimertinib treatment group, which is 8.7 months longer than that in the standard EGFR-TKI group ^[14]^. However, just like any other EGFR-TKIs, Osimertinib inevitably develops acquired resistance, which limits its efficacy in treating patients with EGFR-mutated NSCLC ^[5]^。 The mechanisms of Osimertinib resistance identified so far can be roughly divided into two categories: targeted EGFR-dependent and off-target EGFR-independent mechanisms, including changes in the EGFR pathway, such as the C797S mutation; bypass activation, such as cellular Mesenchymal-Epithelial Transition (cMET) amplification, human epidermal growth factor receptor-2 (HER2) alteration, EGFR amplification, histological type changes, such as conversion to SCLC ^[15]^. The combination of Osimertinib with bypass targeted therapy and other therapeutic approaches is a promising strategy. For example, Anlotinib combined with Osimertinib could reverse acquired Osimertinib resistance in NSCLC by targeting the c-MET/MYC/AXL axis ^[16, 17]^。 At 2022’s American Society of Clinical Oncology (ASCO) meeting, we were very pleased to see the emergence of new drugs to overcome EGFR-TKIs resistance, such as JNJ-372, a bi-specific antibody targeting cMET and EGFR, which was shown to achieve Partial Response (PR) in 25 of 88 evaluated patients, with an effective rate of 28% ^[18]^。

Formononetin is one of the main isoflavone components isolated from Astragalus membranaceus and has been shown to have multiple pharmacological benefits ^[19]^. Formononetin induces cell apoptosis, cell cycle arrest and inhibits cell invasion through the regulating of multiple signaling pathways, including BCL2 associated X (Bax), Bcl-2, and caspase-3, cyclins, signal transduction and transcriptional activator (STAT), Mitogen-Activated Protein Kinase (MAPK) signaling pathways, Phosphatidylinositol 3-Kinase/protein Kinase-B (PI3K/AKT), Vascular Endothelial Growth Factor (VEGF), Fibroblast Growth Factor 2 (FGF2), as well as Matrix Metalloproteinases, plays an important role in the fighting of breast, prostate, and colon cancers ^[20]^. In addition, Combined treatment with other chemotherapeutic agents, such as Bortezomib, LY2940002, U0126, Sunitinib, Epirubicin, Doxorubicin, Temozolomide and Metformin could enhanced the anti-cancer potential of Formononetin through a synergistic effect ^[20]^。 For instance, Formononetin can synergistically enhance the anti-glioblastoma effects of chemotherapeutic drugs ^[21]^. Previously, researchers have discovered that Formononetin can suppress the proliferation of human NSCLC through induction of cell cycle arrest and apoptosis ^[22]^, and inhibits tumor growth by suppressing of EGFR-Akt-Mcl-1 axis in NSCLC ^[23]^. However, the regulatory effects and the potential mechanisms of Formononetin on Osimertinib resistant NSCLC cells were remain unclear.

In this work, we found that Formononetin could significantly induced cell apoptosis but dramatically inhibited cell proliferation and tumor cell clonal formation in Osimertinib resistant NSCLCs. Transcriptome analysis showed tremendous genes expression profiles, and these differentially expressed genes (DEGs) could be clustered into many unique signaling pathways as enriched in GO and KEGG database. Our results not only provided a treatment method to revise the Osimertinib resistance in NSCLC, but also preliminarily revealed the mechanisms from the gene transcriptional levels.

## 2 Materials and Methods

### 2.1 Reagents

The Annexin V-FITC/PI cell apoptosis detection kit (40302ES60) was purchased from Yisheng Biotechnology (Shanghai) Co., LTD. Cell Counting kit-8 (C0038) was the product of Beyotime Biotechnology Co., LTD. TRIzol^TM^ RNA extraction reagent (15596018) and CloneMinerTM □cDNA library construction reagent (A11180) was purchased from ThermoFisher Scientific. Osimertinib (HY-15772) and Formononetin (HY-N0183) was the product from MedCheExpress (MCE).

### 2.2 Cell culture

H1975 (EGFR L858R/T790M) NSCLC cell line was purchased from Wuhan Pricella Biotechnology Co., LTD. and was maintained in Roswell Park Memorial Institute (RPMI-1640) growth medium supplemented with 10% fetal bovine serum (FBS, Invitrogen) and 1% penicillin-streptomycin (Invitrogen) at 37°C in the presence of 5% CO_2_. The cells were sub-cultured every two days upon reaching 90% confluency.

### 2.3 Generation of Osimertinib-resistant cell lines

Osimertinib-resistant H1975 cell line was established as described previously ^[24]^. In brief, approximately 1 × 10^6^ H1975 cells were seeded in 10 cm cell culture dish. 500□nM Osimertinib was used to treat the cells. Medium changed every 2 days and the exposure dose was increased by 500□nM every 15 days until achieving the final concentration of 2 µM. Osimertinib treated cells were maintained for 2 months and heterogenous H1975 Osimertinib-treated H1975 cells were further seeded into a 96-well plate using limiting dilution. All clones were maintained in RPMI-1640 complete medium supplemented with 4 µM Osimertinib for 4 months. Finally, the remaining cell clones were selected as Osimertinib-resistant H1975 cell lines, and were named as H1975_OR.

### 2.4 Cell apoptosis assay

1 × 10^5^ H1975_OR cells per well were seeded into a 6-well cell culture plate and cultured overnight at 37□ in the presence of 5% CO_2_. A final concentration of 1.25 μM, 2.5 μM and 5 μM Formononetin was added into each well of the cell, respectively. For control cells, add same volume of solvent. 24h later, the cells were digested using 0.2 mL of 0.25% EDTA-free trypsin for 3 min. The cells were washed twice with pre-cooled PBS and centrifuged at 4□ 300 g for 5 min. PBS was absorbed and 100 μL of 1 × binding buffer was added to re-suspension cells. 5 μL of Annexin V and 10 μL PI staining solution per sample were added and lightly mixed. The samples were incubated protected from light, under room temperature for 15 mins. After diluted 5 times using 1 × binding buffer, the cells were immediately tested using a BD FACSCelesta^TM^ flow cytometry.

### 2.5 Cell proliferation assay

1 × 10^3^ H1975_OR cells per well were seeded into a 96-well cell culture plate and cultured overnight at 37□ in the presence of 5% CO_2_. A final concentration of 1.25 μM, 2.5 μM and 5 μM Formononetin was added into each well of the cell. Then on 0h, 24h, 48h, 72h and 96h after Formononetin treatment, 10 μL of Cell Counting Kit-8 was added into the cell culture medium and incubated for 4h and the absorbance at 450 nm were tested using a spectrophotometer. Control cells were treated with same volume of solvent.

### 2.6 Clonal formation assay

5 × 10^2^ H1975_OR cells per well were seeded into a 6-well cell culture plate and cultured overnight at 37□ in the presence of 5% CO_2_. A final concentration of 1.25 μM of Formononetin was added into each well. For control cells, add same volume of solvent. The cells were cultured for 10 days at 37□ in the presence of 5% CO_2_ to form cell clones. The cell culture medium was changed for fresh complete medium every 3 days. Each time after the medium was changed, Formononetin with a final concentration of 1.25 μM was added to the Formononetin treatment cells. Control cells were added with same volume of solvent. After cell clone was formed, the cells were washed twice with PBS, and 1 mL of 4% paraformaldehyde was added to each well for 30 mins, after washed with PBS, add 1 mL of crystal violet dye solution to each well and kept for 20 min to stain the cells. After washed with PBS several times, the cell clones were dried, photographed and counted.

### 2.7 Cell treatment and total RNA extraction

2 × 10^6^ of H1975_OR cells were seeded into a 10 cm cell culture dish and were cultured overnight at 37□ in the presence of 5% CO_2_. A final concentration of 2.5 μM Formononetin was added into the cell culture medium. Cells in control group were treated with same volume of solvent. 24h later, the cell medium was removed and the cells were washed twice with pre-cooled PBS. Then 1 mL of TRIZOL reagent was added to each well, shaken well, and left at room temperature for 5 min to lyse the cells. The cell lysate was taken into a 1.5 mL RNase-free Eppendorf (EP) tube, and 0.2 mL chloroform was added, and gently shaken for 15 s. After incubated at room temperature for 5 min, centrifuged at 4□, 12000 rpm for 15 min. Then take the upper water phase (about 0.5 mL) into a new 1.5 mL RNase-free EP tube, add 0.5 mL isopropyl alcohol, mix well, and incubated at room temperature for 10 min. Centrifuged at 4□, 12000 rpm for 10 min. Discard the supernatant and washed twice with 1 mL 75% ethanol. After dried in air for 5 min, add 50 μL of DEPC-treated water and hold in the water bath at 55□ for 10 min to completely dissolve total RNA, and then the concentrations were measured. The total RNA extracted was tested for purity, quantity, and integrity.

### 2.8 Library construction and sequencing

Libraries were constructed according to the protocols supplied by the manufacture. In brief, 1 μg of total RNA per sample were used to generate the library. Oligo dT magnetic beads was used to bind specifically to the poly (A) tail of mRNA to remove other RNAs. And then the mRNAs were fragmentated using the fragmentation reagent. Using a random hexamer primer, reverse transcriptase to synthesizes first strand cDNA using mRNA as a template and the synthesizes a second strand, and produces a double-stranded cDNA (ds cDNA). After end repair of double-stranded cDNA, end complement, and then the repaired cDNAs were purified. Finally, the libraries were enriched by polymerase chain reaction (PCR). After amplified and purified, the libraries were ready for sequencing. The RNAs sequencing were performed on an Illumina NextSeq 2000 platform.

### 2.9 Gene Ontology (GO) Enrichment analysis

TopGO R package was used for GO enrichment analysis. During the analysis, differential genes annotated by GO term were used to calculate the gene list and gene number of each term, and then *p* value was calculated by hypergeometric distribution method (the threshold for significant enrichment was set as *p* value < 0.05) to find out that, compared with the whole genome background, significant enrichment of differential genes in GO term. According to the GO enrichment results, the enrichment degree was measured by Rich factor, False discovery rate (FDR) value and the number of genes enriched to GO terms. Select the top 20 GO term entries with the smallest FDR value for presentation.

### 2.10 Kyoto Encyclopedia of Genes and Genomes (KEGG) Enrichment analysis

ClusterProfiler R package was used for KEGG enrichment analysis with the *p* cutoff value set as < 0.05. According to the results of KEGG enrichment, the enrichment degree was measured by Rich factor, FDR value and the number of genes enriched on this pathway. The top 20 KEGG pathways with the smallest FDR value were selected for display.

### 2.11 Gene Set Enrichment Analysis (GSEA) Enrichment analysis

GSEA is a computational method used to determine whether a priori defined gene set shows a statistically significant, consistent difference between two biological states. We use GSEA software to generate the enrichment score (ES). Normalized Enrichment Score (NES) is obtained by standardizing ES calculated for each gene subset according to the size of the gene set. The FDR is then calculated for NES. Pathways with |NES| > 1, *p* value < 0.05 and FDR < 0.25 were considered significantly enriched.

### 2.12 Data availability

The datasets produced in this study are available in the following databases: RNA-seq data: NCBI SRA: PRJNA687994 (https://www. ncbi.nlm.nih.gov/bioproject/PRJNA687994).

### 2.23 Statistical analysis

Data are shown as the mean ± SD. and performed using Prism 5 (GraphPad Software). Statistical significance between two experimental groups was calculated using unpaired two-tailed Student’s *t* test. Multiple sets of data comparison were calculated by analysis of variance (ANOVA). Cell proliferation assay were tested using a two-tailed Mann–Whitney *U* test. *, p < 0.05, **, p < 0.01, ***, p < 0.001, NS, Non-significant, *p* > 0.05.

## 3 Results

### 3.1 Formononetin significantly promoted cell apoptosis, inhibited cell proliferation and clonal formation in H1975_OR cells

Previous works have shown that Formononetin could suppressed the proliferation and induced cell cycle arrest and apoptosis in human NSCLC ^[22, 23]^. In this work, we set out to investigate the potential pharmacological effects of Formononetin in the regulation of the tumor biological process in Osimertinib-resistant NSCLC cells. We firstly treated Osimertinib-resistant NSCLC cell line H1975_OR with 1.25 μM, 2.5 μM and 5 μM Formononetin, and 24h later, cell apoptosis was analyzed using flow cytometry. It can be seen in **Figure 1A-C** that Formononetin could dose-dependently induced both early (Annexin V^+^PI^-^) and late cell apoptosis (Annexin V^+^/PI^+^) in H1975_OR cells as compared to solvent controls. 1.25 μM and 2.5 μM of Formononetin slightly induced H1975_OR cell growth, but the growth rate was significantly lower than in solvent controls (**Figure 1D**). Whereas, the cell viabilities in 5 μM Formononetin-treatment group were significantly inhibited (**Figure 1D**). In addition, Formononetin also significantly inhibited clonal formation in H1975_OR cells as compared to negative controls (**Figure 1E and F**). These results indicated that Formononetin could significantly induced cell apoptosis, inhibited cell proliferation and tumor clonal formation in H1975_OR cells.

**Figure 1.**
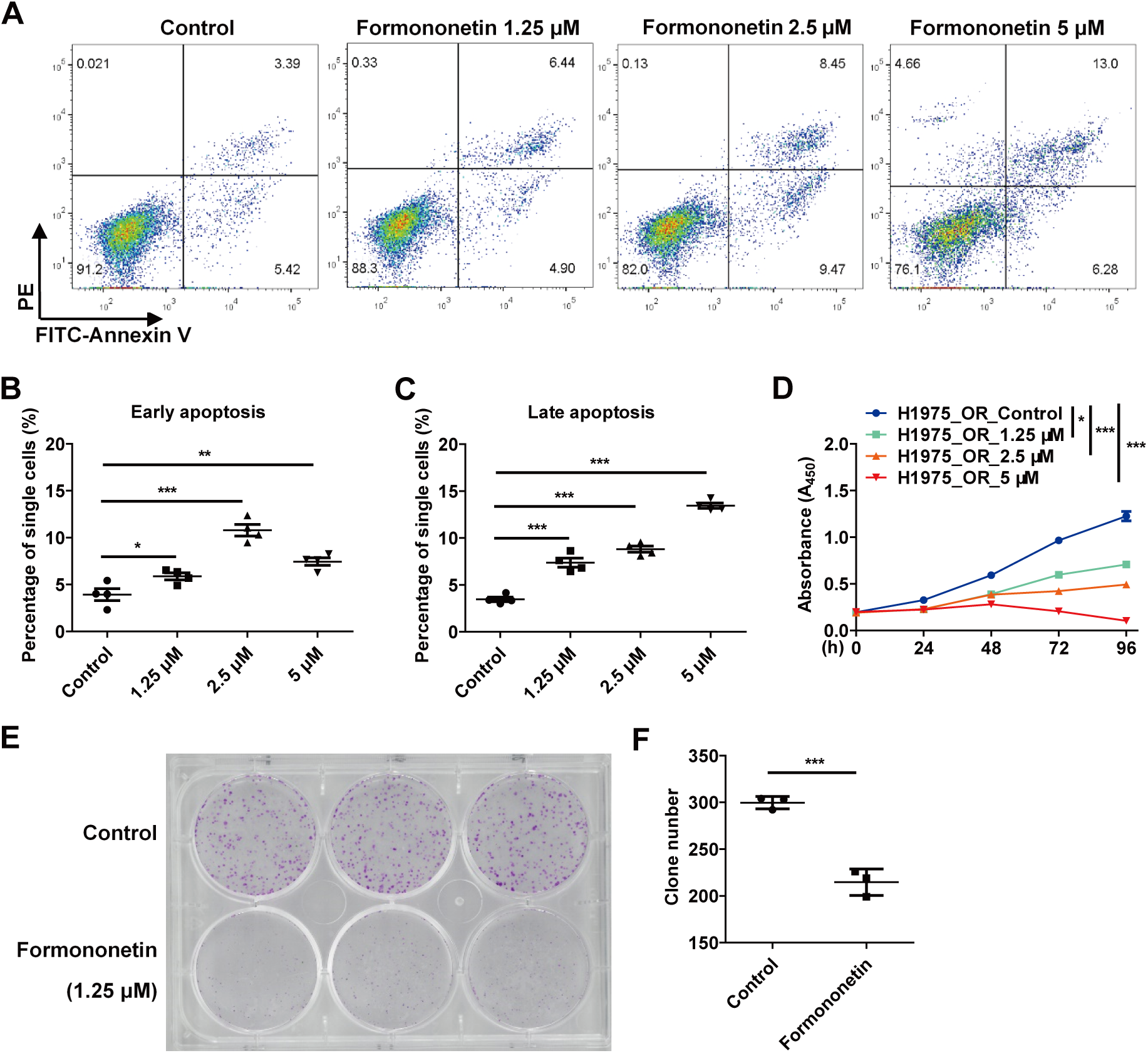
**Formononetin promoted cell apoptosis, inhibited cell proliferation and clonal formation in H1975 Osimertinib resistant lung cancer cells.** (**A-C**) H1975 Osimertinib resistance cells were treated with a serial dose of Formononetin from 1.25 to 5 μM for 24h, and cell apoptosis was tested by Flow cytometry (**A**). The early apoptosis cells (Annexin V positive, PI negative, Annexin V^+^PI^-^) (**B**) and the late apoptosis cells (Annexin V positive, PI positive, Annexin V^+^PI^+^) (**C**) were calculated. (**D**) Cell proliferation assay. H1975 Osimertinib resistance cells were treated with a serial dose of Formononetin from 1.25 to 5 μM for 0h, 24h, 48h, 72h and 96h and cell proliferation was tested by Cell Counting kit-8 (CCK-8) method. (**E**) A representative picture of clonal formation. H1975 Osimertinib resistance cells was treated with 1.25 μM Formononetin for 14 days, and the cell clones were fixed with 4% paraformaldehyde and stained with crystal violet. (**F**) Number of the clones. Data are shown as the mean ± SD., and are representative of three independent experiments. Significant differences were tested using analysis of variance (ANOVA) (**B, C**), two-tailed Mann–Whitney *U* test (**D**) and two tailed student *t* text (**F**). P < 0.01, **P < 0.01, ***P < 0.001.

### 3.2 Figure 2 Formononetin significantly altered the gene expression profiles in H1975_OR cells

To study the potential mechanisms for Formononetin counteracted-Osimertinib resistance in NSCLC, we firstly conducted a transcriptome sequencing analysis of the Formononetin treated H1975_OR cells and solvent controls. Principal Component Analysis (PCA) showed good intergroup difference and intragroup consistency between Formononetin treated H1975_OR cells and solvent controls (**Figure 2A**). Formononetin induced a vast of Differentially Expressed Genes (DEGs) in H1975_OR cells as compared to solvent controls (**Figure 2B and C**). Concretely speaking, Formononetin significantly promoted 2213 genes expression but dramatically inhibited 2096 genes expression in in H1975_OR cells as compared to solvent controls (**Figure 2D**). These results indicated that Formononetin significantly altered the gene expression profiles in H1975_OR cells.

**Figure 2.**
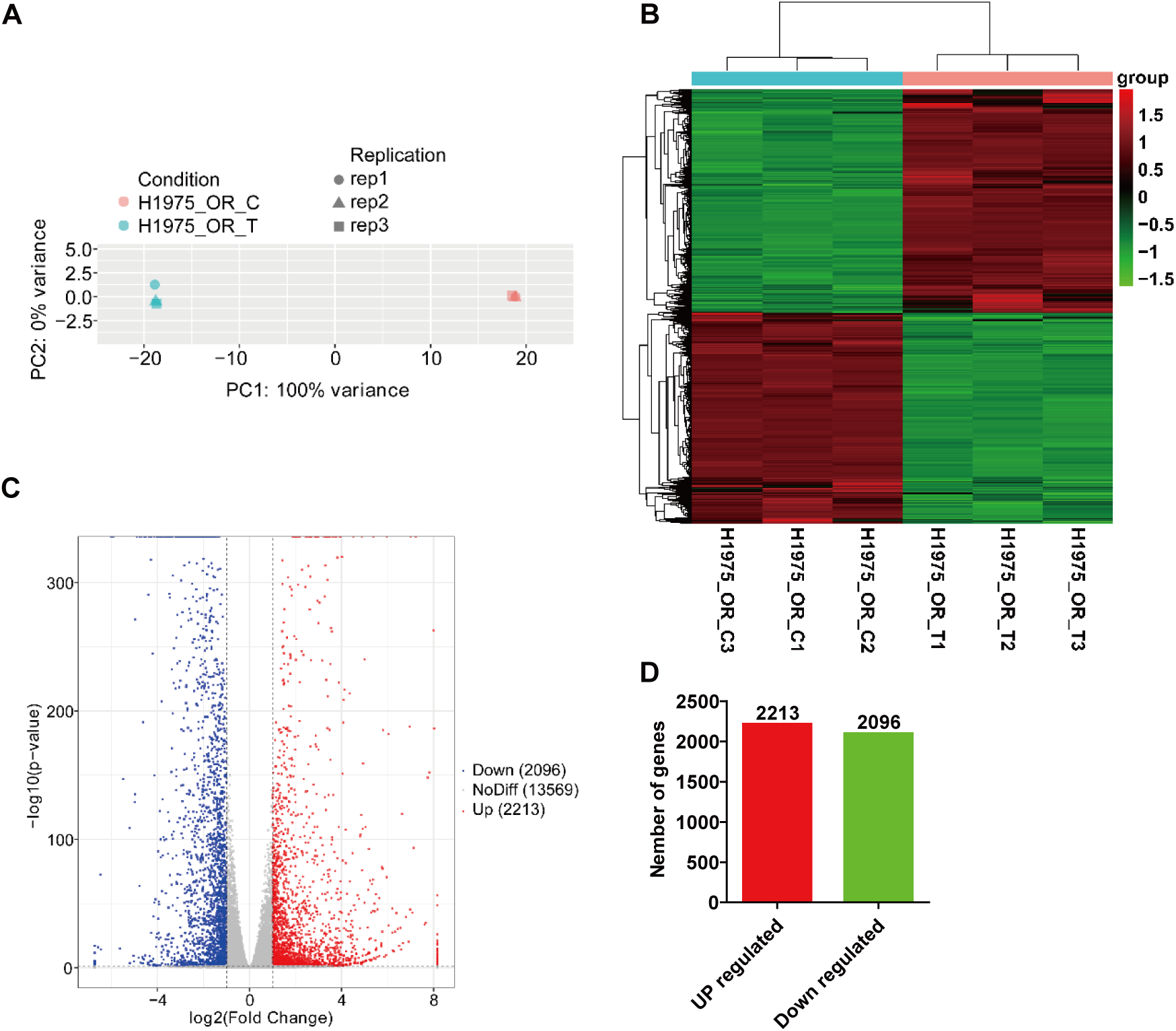
**Formononetin significantly altered the gene expression profiles in Osimertinib resistant lung cancer cells.** H1975 Osimertinib resistant lung cancer cells were treated with or without 1.25 μM Formononetin for 24h, the mRNA from the samples were extracted and sequenced on an Illumina NesxtSeq 2000 secondary sequencing platform. (**A**) Principal component analysis (PCA) of the samples. (**B**) Heatmap shows the differentially expressed genes (DGE). (**C**) Volcano plot shows the DEGs. (**D**) Dis-regulated gene numbers between Formononetin treated and nontreated cells.

### 3.3 Figure 3 Gene Ontology (GO) and Kyoto Encyclopedia of Genes and Genomes (KEGG) analysis

GO is a representation of the nature of a body of knowledge in a biological domain, usually consisting of a set of classes (or terms or concepts) with relationships between them and describes what we know about the field of biology in terms of GO domains: molecular functions, cellular components, and biological processes ^[25]^. In this work, we found that Formononetin induced DEGs in H1975_OR cells can be clustered into a vast of pathways in GO database, among which, Anatomical structure morphogenesis, Anatomical structure development, Multicellular organism development and Developmental process to be the top 4 most significantly altered pathways according to the enrichment rich factor and FDR values (**Figure 3A**). Furthermore, **Figure 3B** shows TOP 20 most significantly altered KEGG pathways in Formononetin treated H1975_OR cells, among which, MAPK, Axon guidance, pathway in cancer, Amoebiasis and AGE-RAGE signaling pathway in diabetic complications to be the most dramatically altered KEGG pathways in H1975_OR cells upon Formononetin treatment.

**Figure 3.**
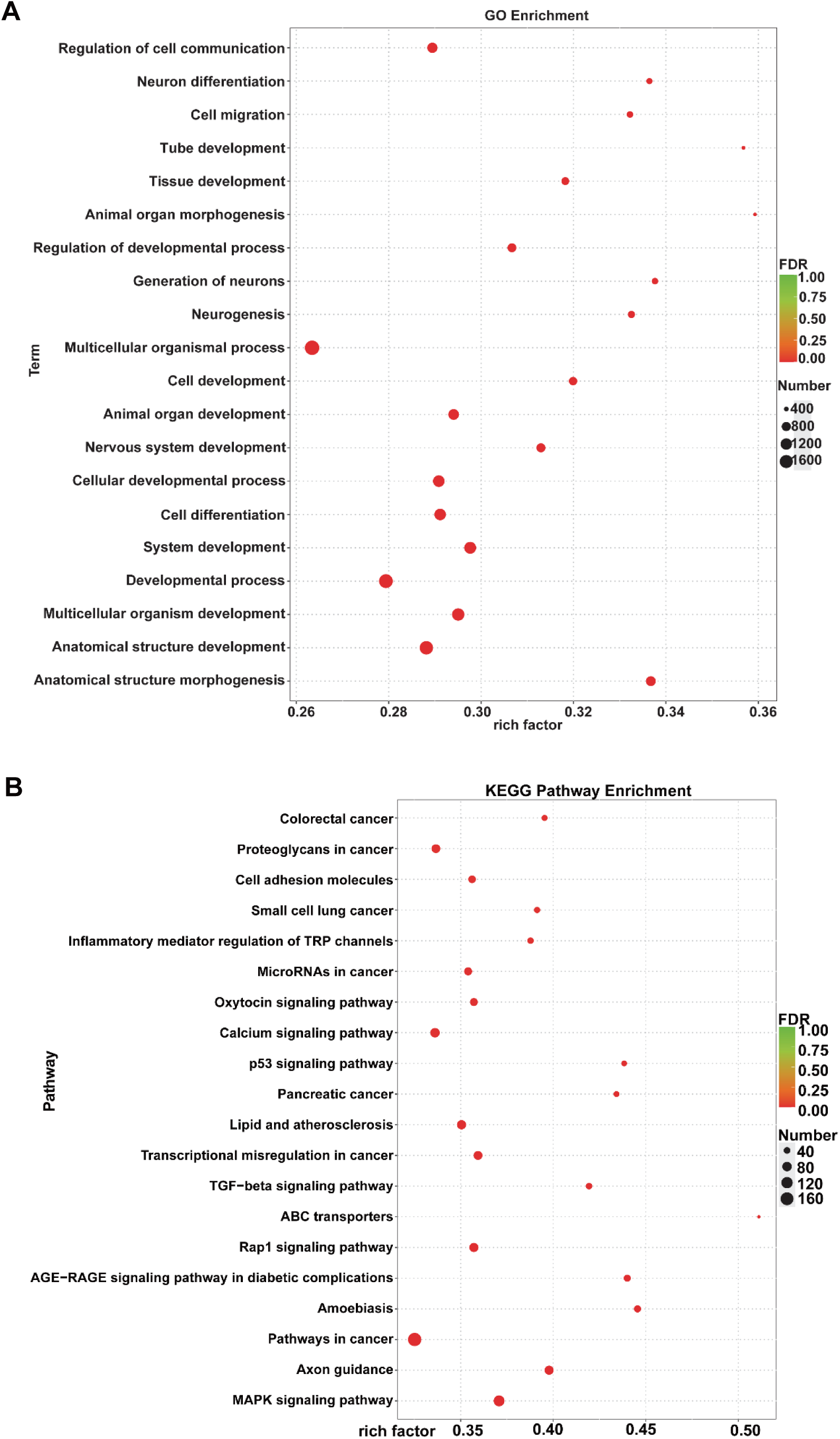
**Gene Ontology (GO) and Kyoto Encyclopedia of Genes and Genomes (KEGG) analysis.** (**A**) Bubble chart shows top 20 significantly dis-regulated pathways in GO enrichment. (**B**) Bubble chart shows top 20 significantly dis-regulated pathways in KEGG pathway enrichment.

### 3.4 Figure 4 Formononetin significantly regulated genes expression associated to inflammatory signaling pathways in Osimertinib resistant lung cancer cells

4 most significantly altered inflammatory signaling pathways of Formononetin treated Osimertinib resistant lung cancer cells were shown in **Figure 4**. **Figure 4A** shows top 10 most significantly up-regulated and top 10 most dramatically down-regulated genes in MAPK signaling pathway, among which, PGF, HSPA6, RASGRP4 and CACNA1A to be the mostly promoted and ANGPT1, PRKCA, PRKCB and CACNA2D1 to be the mostly inhibited genes in response to Formononetin treatment. In addition, in TGF-beta signaling pathway, RGMA, FST, NOG and INHBA to be the most significantly up-regulated and BMP6, BMPR1B, THSD4 and LTBP1 to be the mostly down-regulated genes upon Formononetin treatment (**Figure 4B**). What’s more, Formononetin treatment significantly inhibited GTSE1, COP1, ATM and RRM2 expression but dramatically promoted SERPINB5, RPRM, CD82 and GADD45G expression in Osimertinib resistant lung cancer cells, all of which were associated to P53 signaling pathway (**Figure 4C**). Formononetin significantly promoted IL36G, CCL22, LTA and NGFR expression but dramatically inhibited CCL8, CX3CR1, CCR1 and BMP6 expression in Osimertinib resistant lung cancer cells, all of which were associated to Cytokine-cytokine receptor interaction signaling pathway (**Figure 4D**). These results indicated that Formononetin could significantly regulated genes expression associated to inflammatory signaling pathways in Osimertinib resistant lung cancer cells.

**Figure 4.**
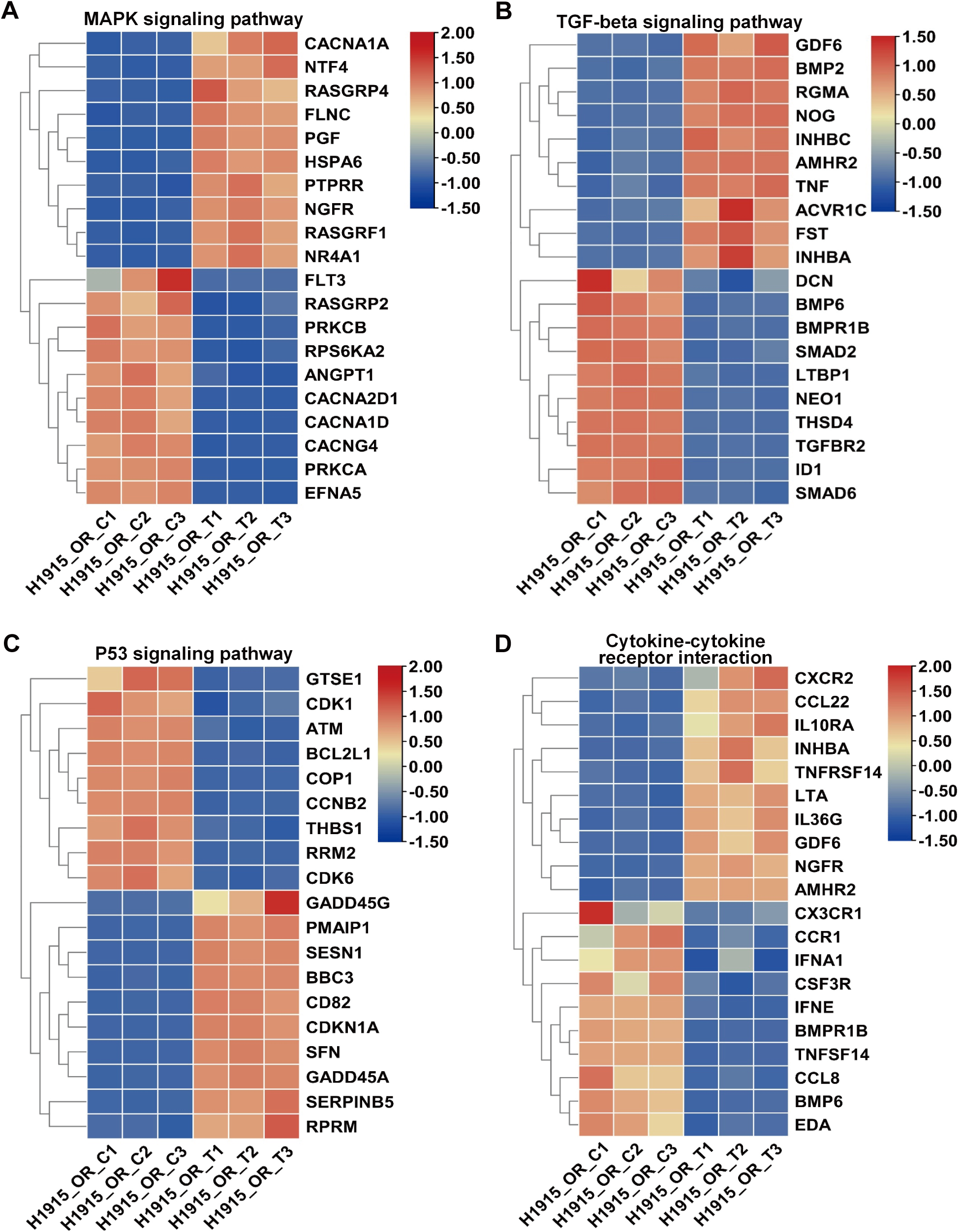
**Formononetin significantly regulated genes expression associated to inflammatory signaling pathways in Osimertinib resistant lung cancer cells.** (**A**) Heatmap shows top 10 most significantly up-regulated and top 10 most significantly down-regulated genes in TGF-beta signaling pathways. (**B**) Heatmap shows top 10 most significantly up-regulated and top 10 most significantly down-regulated genes in MAPK signaling pathways. (**C**) Heatmap shows top 10 most significantly up-regulated and 9 significantly down-regulated genes in P53 signaling pathways. (**D**) Heatmap shows top 10 most significantly up-regulated and top 10 most significantly down-regulated genes in Cytokine0cytokine receptor interaction signaling pathways.

### 3.5 Formononetin significantly regulated genes expression associated to metabolism in Osimertinib resistant lung cancer cells

Figure 5 shows 4 most significantly dis-regulated pathways associated to metabolism. Formononetin significantly inhibited CYP2A7, CYBB, PRKCA and PLCB4 genes expression but significantly promoted SELE, HSPA6, POU2F3 and CYP1A1 genes expression, which were associated to Lipid and atherosclerosis pathway (Figure 5A). In addition, in Purine metabolism signaling pathway, PDE7B, GUCY1A2, PDE11A and RRM2 to be the most significantly inhibited genes and PDE4C, PDE2A, XDH and PDE6G to be the most significantly promoted genes upon Formononetin treatment (Figure 5B). What’s more, Formononetin significantly up-regulated CHST4, CHST1, CHST6, CHST2 and B3GNT7 expression, but significantly down-regulated FUT8, ST3GAL1 and B4GALT1 expression in Glycosaminoglycan biosynthesis-keratin sulfate pathway upon Formononetin treatment (Figure 5C). Formononetin significantly promoted NAT1 and XDH expression but dramatically inhibited CYP2A7 and CYP2A6 in Osimertinib resistant lung cancer cells, all of which were associated to Caffeine metabolism (Figure 5D). These results indicated that Formononetin significantly regulated genes expression associated to metabolism in Osimertinib resistant lung cancer cells.

**Figure 5.**
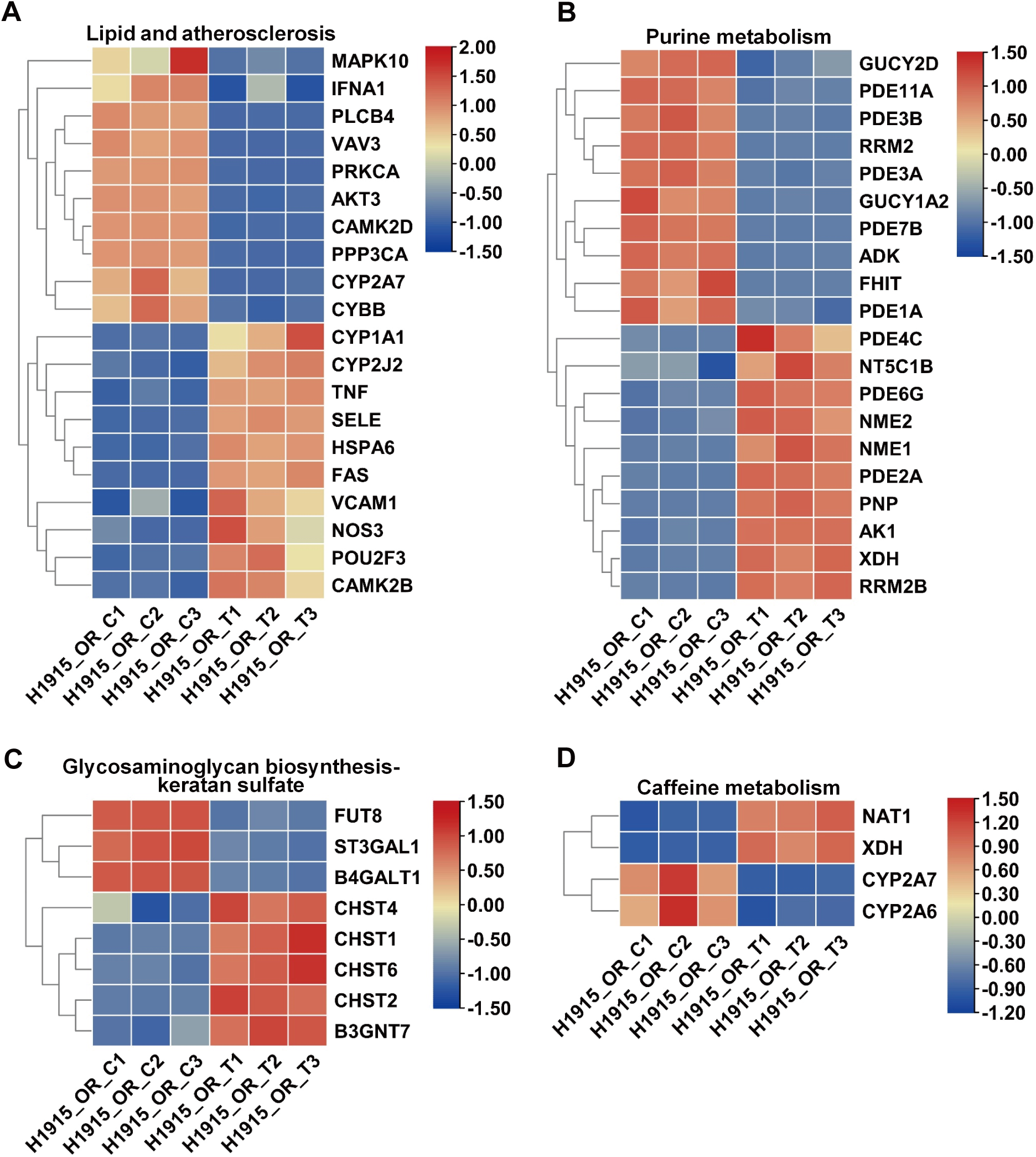
**Formononetin significantly regulated genes expression associated to metabolism in Osimertinib resistant lung cancer cells.** (**A**) Heatmap shows top 10 most significantly up-regulated and 10 most significantly down-regulated genes associated to Lipid and atherosclerosis signaling pathways. (**B**) Heatmap shows top 10 most significantly up-regulated and 10 most significantly down-regulated genes associated to Purine-metabolism signaling pathways. (**C**) Heatmap shows DEGs associated to Glycosaminoglycan biosynthesis-keratan sulfate signaling pathways. (**D**) Heatmap shows DEGs associated to Caffeine metabolism signaling pathways.

### 3.6 GSEA analysis

In order to explore the potential mechanisms of Formononetin counteracted Osimertinib resistant in lung cancer cells, the GSEA analysis was used to predict the KEGG downstream pathway. Figure 6 show top 4 mostly dysregulated pathway in GSEA analysis. The result showed that Formononetin could activated the hallmarks of Allograft rejection and Autoimmune thyroid disease (**Figure A and B**), but inhibited hallmarks of Aldosterone-regulated sodium reabsorption and Bacterial invasion of epithelial cells (**Figure C and D**). These results indicated that Formononetin could significantly regulate the state of the signaling pathways in KEGG enrichment.

**Figure 6.**
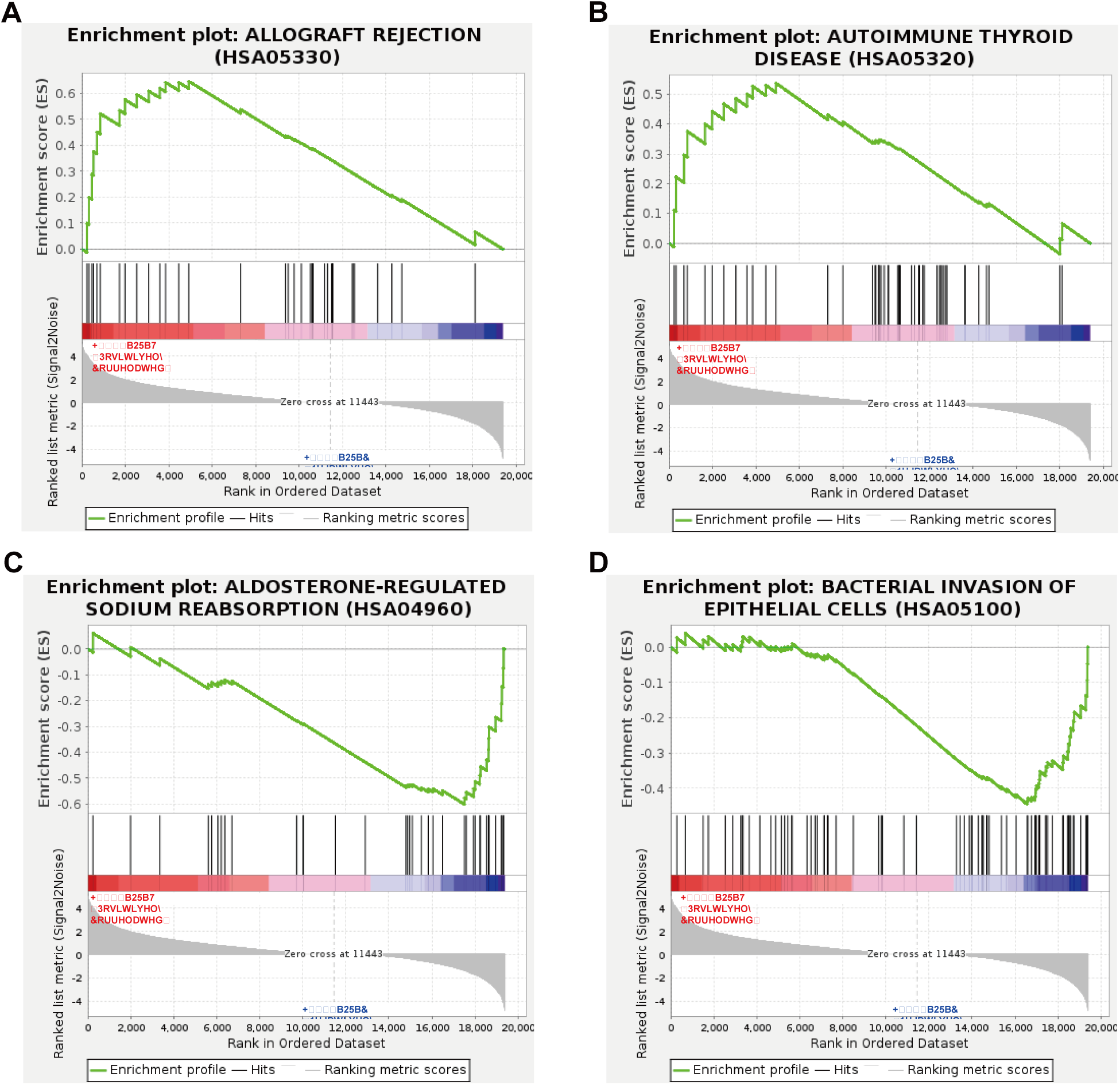
GSEA analysis. (**A**) GSEA analysis of genesets for Allograft rejection. (**B**) GSEA analysis of genesets for Autoimmune thyroid disease. (**C**) GSEA analysis of genesets for Aldosterone-regulated sodium reabsorption. (**D**) GSEA analysis of genesets for Bacterial invasion of epithelial cells. NES, Normalized enrichment score. FDR, False discovery rate. Positive and Negative NES indicate higher and lower expression in Formononetin treated and nontreated H1975 Osimertinib resistance lung cancer cells.

## Discussion

EGFR gene mutation is very common in lung cancer patients in China, with 51.4% advanced lung adenocarcinoma patients with EGFR sensitive mutations, whereas, in non-smoking adenocarcinoma patients, the mutation rate is as high as 60% ^[26]^. The role of the EGFR gene is to control the cell’s synthesis of the EGFR, which binds to the EGF in body fluids, thus initiating a series of biochemical reactions that promote the synthesis of DNA and mitosis. When a cell’s EGFR gene is mutated, the EGFR produced by the cell continues to send a “division signal” to the cell, causing the cell to proliferate without control and evade apoptosis, which eventually result in a malignant tumor ^[5]^. To treat this type of lung cancer, scientists have created three generations of drugs that specifically target different types of mutations and bind to EGFR on cancer cells, thereby controlling tumor growth, preventing tumor angiogenesis and inducing cancer cell death ^[17]^. Prior to the birth of Osimertinib, the usage of first-and second-generation EGFR-TKIs (Gefitinib, Erlotinib, Afatinib, etc.) had extended progression-free survival (PFS) in patients of NSCLC with EGFR mutations (except for exon 20 insertion mutations) from 5 months to more than 9 months, which is equivalent of chemotherapy could achieve ^[27]^. Although the emergence of EGFR-TKIs has brought revolutionary breakthroughs to the treatment of lung cancer, the disease will still progress in 9 - 11 months after treatment with EGFR-TKIs ^[28]^. More than 60% of these patients have a new T790M mutation in the EGFR gene. This mutation further alters the structure of EGFR, causing patients to become resistant to first - or second -generation of EGFR-TKIs ^[29]^. Osimertinib, the third generation of EGFR-TKIs, is an irreversible EGFR inhibitor that inhibits cancer progression caused by EGFR-TKIs sensitive and T790M resistant mutations, as well as exon 19 and 21 mutations ^[30]^. Osimertinib was shown to have a mPFS of 18.9 months, compared with 10.2 months for Gefitinib and Erlotinib: the life expectancy of patients with EGFR mutant lung cancer has been extended by 8 months again ^[12]^.

The emergence of Osimertinib has largely solved the problems of first-and second-generation drug resistance in lung cancer patients. Unfortunately, however, patients still develop resistance again after using Osimertinib for a period of time. At present, a series of resistance mechanisms of Osimertinib have been discovered, mainly including: 1. The EGFR gene is mutated again. Osimertinib overcomes resistance arising from the T790M mutation by irreversibly binding to the EGFR-C797 site ^[30]^. After a period of medication, the secondary EGFR gene C797S mutation will develop drug resistance again. In addition, other EGFR site mutations may occur during treatment, including EGFR-L718Q, -L844V, -L798I, -L692V, -E709K, etc ^[30]^. 2. Drug resistance mechanisms associated with other bypass pathways. In particular, after the T790M mutant clone was reduced or even completely eliminated, tumor cells continued to multiply, indicating that they had other resistance mechanisms or relied on other pathways other than EGFR ^[15]^. It may include HER2 amplification ^[31]^, MET amplification ^[32]^, Hepatocyte Growth Factor (HGF) expression ^[33]^, PIK3CA mutation ^[34]^, PTEN deletion ^[35]^, KRAS mutation ^[36]^, IGF1R activation ^[37]^ and gene fusion ^[38]^, etc. 3. Disease resistance occurs as a result of the transformation from NSCLC to SCLC ^[15]^. Considering that there were so many resistance mechanisms, each of which is enough to trigger Osimertinib resistance, and that there were a variety of different combinations of resistance mechanisms in different patients, which undoubtedly greatly increases the difficulty of scientific research ^[15]^.

Osimertinib resistant patterns are divided into local progression, slow progression, and systemic or fulminant progression ^[15]^. Different resistance patterns require different coping strategies. Osimertinib or combination of radiotherapy, surgery, or antiangiogenic targeting agents is recommended for patients with local or slow progression. For more than 50% of patients with slow disease progression, the mPFS can be extended by 3 - 5 months ^[15]^. For patients with fulminant progression, the relationship between the C797S mutation and the EGFR-T790M mutation must first be clarified ^[39]^. If the patient is resistant to first, second, and third generation EGFR-TKIs, and there is still lacking of mature treatment options. Second-generation ALK-TKI Brigatinib can competitively bind to the ATP binding site of the EGFR kinase domain and is not affected by T790M or C797S steric hindrance, so it can inhibit the growth of L858R and T790M and C797S triple mutation tumors and delay Osimertinib-induced acquired resistance ^[40]^.

Combination therapy is an effective method to delay Osimertinib resistance. For instance, Osimertinib combined with Savolitinib have been reported to break through this drug resistance bottleneck, in which, all lung cancer patients experienced disease progression after treatment with Osimertinib, among whom, 62% of patients with MET overexpression and/or amplification, and more than one-third (34%) had MET hyperexpression ^[41]^. The Objective Partial Response (OPR) of Osimertinib combined with Savolitinib was as high as 49% in patients with high levels of MET expression ^[42]^. Furthermore, recent studies have shown that small molecules in traditional Chinese medicine have the potential to reverse Osimertinib resistance. Cai and his colleagues found that Dihydroartemisinin can inhibits NSCLC resistant cell proliferation by downregulating the expression of heme oxygenase 1 (HO-1) and inhibiting Osimertinib resistant EGFR mutations ^[43]^. In this paper, we found that the anti-NSCLC drug Formononetin can dose-dependently induce apoptosis, cell proliferation, and clonal formation of Osimertinib-resistant NSCLC cells, which may be achieved by regulating the immune function status and metabolic reprogramming process of Osimertinib-resistant cells.

Formononetin belongs to the isoflavones and is mainly found in leguminous plants such as the roots of astragalus ^[22]^. In recent years, researchers have done a lot of work on the pharmacological effects of Formononetin, showing that it has anti-inflammatory, antioxidant, anti-apoptotic, lipid-modulating, antithrombotic, tumor inhibition effects ^[19]^. In addition, Formononetin can also reverse multidrug resistance in Adriamycin-resistant hormone receptor-positive (MCF-7/ADR) breast cancer cells, possibly by inhibition of autophagy and reduction of P-glycoprotein (P-gp) protein ^[20]^. In addition, studies have found that Formononetin can reduce the deterioration of triple-negative breast cancer (TNBC) by inhibiting lncRNA AFAP1-AS1-miR-195/miR-545 axis and can also improve drug sensitivity of paclitaxel-resistant TNBC by inhibiting autophagy ^[44]^. Through the *in vitro* investigation of metabolism of rat intestinal flora and the usage of cell membrane fixation chromatography technology, Yang and his colleagues found that Formononetin may be an effective component of astragalus in the treatment of lung cancer, and they may play a role in the treatment of lung cancer through the induction of autophagy and p53/AMPK/mTOR signaling pathways ^[45]^. In lung cancer, Formononetin suppresses the proliferation of human NSCLC cells through induction of cell cycle arrest and apoptosis ^[22]^. Yu et al. screened 98 commercially available natural products and found that Formononetin could inhibits both WT and mutant EGFR kinase activity *in vitro* and *in vivo* ^[23]^. Formononetin inhibits EGFR-Akt signaling, thereby activating GSK3β and promoting Mcl-1 phosphorylation and ubiquitination and degradation in NSCLC cells, suggesting the importance of promoting ubiquitination-dependent Mcl-1 switching may be an alternative strategy to enhance the anti-tumor efficacy of EGFR-TKI ^[23]^. However, the potential functions of Formononetin on the regulation of Osimertinib resistance in NSCLC cells were largely unknown. In this work, we found that Formononetin can significantly induced Osimertinib resistant cells apoptosis, proliferation and inhibited clonal formation of Osimertinib resistant cells.

In conclusion, in this work, we found that Formononetin could significantly induced cell apoptosis but dramatically inhibited cell proliferation and clonal formation in Osimertinib resistant NSCLCs. Our results not only uncovered the underlying mechanisms for Formononetin to reverse NSCLC resistance Osimertinib, but also provided a potential treatment method for Osimertinib resistant NSCLC patients.

## Data availability

The datasets produced in this study are available in the following databases: NCBI BioProject with the accession number: PRJNA1021017.

## CrediT Contribution Statement

Conceptualization Yang Liu, Runyang Yin and Jing Gao; Formal Analysis: Runyang Yin and Jing Gao; Writing original draft: Runyang Yin, Jing Gao and Yang Liu.

